# Stronger population suppression by gene drive targeting *doublesex* from dominant female-sterile resistance alleles

**DOI:** 10.1101/2025.04.16.649134

**Authors:** Weizhe Chen, Ziye Wang, Jackson Champer

## Abstract

CRISPR homing drives can be used to suppress a population by targeting female fertility genes. They convert wild-type alleles to drive alleles in the germline of drive heterozygotes by homology-directed repair after DNA cleavage. However, resistance alleles produced by end-joining pose a great threat to homing drive. They prevent further recognition by Cas9, and therefore weaken suppressive power, or even stop suppression if they preserve the function of the target gene. We used multiplexed gRNAs targeting *doublesex* in *Drosophila* to avoid functional resistance and create resistance alleles that were dominant female-sterile. This occurred because the male *dsx* transcript was generated in females by disruption of the female-specific splicing acceptor site. We rescued dominant sterility of the drive by providing an alternate splicing site. As desired, the drive was recessive female sterile and yielded high drive inheritance among the progeny of both male and female drive heterozygotes. The dominant-sterile resistance alleles enabled stronger suppression in computational models, even in the face of modest drive efficiency and fitness costs. However, we found that male drive homozygotes were also sterile because they used the rescue splice site. Attempts to rescue males with alternate expression arrangements were not successful, though some male homozygotes had less severe intersex phenotypes. Though this negatively impacted the drive, models showed that it still had significantly improved suppressive power. Therefore, this design may have wide applicability to *dsx*-based suppression gene drives in a variety of organisms with intermediate homing drive performance.

## Introduction

Gene drive technology has the potential to address some of the most pressing challenges in public health, agriculture, and conservation. It holds significant promise for efficient and environmentally friendly insect control, especially with the rise of gene editing technologies^1–7^.

Gene drives are selfish genetic elements that can bias their own inheritance among progeny at a super-Mendelian frequency (>50%). This makes them capable of spreading desirable traits throughout wild populations or even directly suppressing target populations^1,2^. CRISPR-Cas9 based homing drives, one of most well-studied drive systems, can pass the drive allele on to their offspring at an increased rate by cleaving the wild-type allele in the germline of heterozygotes using a guide RNA (gRNA). The DNA is then repaired by homology-directed repair, using the drive allele as a template. Thus, wild-type alleles are converted into drive alleles in germline cells, increasing drive allele inheritance among the progeny^1,4,5,8^.

Though the drive allele can spread over the whole population in theory, a substantial obstacle is caused by end-joining repair, which cannot be fully avoided in CRISPR-based gene drive^6,9,13,14^. End-joining repair at the target site can induce formation of resistance alleles that prevent the gRNA from recognizing the target. Functional resistance alleles (in which the target gene still works correctly) can rapidly accumulate in the population and stop the drive^9–13^. Nonfunctional resistance alleles that are resistant to cleavage but lose target gene function can slow the spread of gene drive^14^. Resistance remains one of the key challenges of developing efficient CRISPR homing drive approaches. Strategies such as using multiplexed gRNAs and targeting highly conserved regions of genes associated with viability or fertility have proven effective in mitigating the generation of resistance^11,12,15,16^.

For suppression drive systems, an important characteristic is their power to strongly suppress and eliminate target populations. With imperfections, a suppression drive will often reach an equilibrium rather than 100% frequency, thus leaving some fertile individuals. Genetic load is an indicator for evaluating the suppressive power of pest control strategies. It is specifically defined in the context of gene drive as the average net fitness reduction relative to a wild-type population of the same size after the drive reaches its equilibrium. In short, high genetic load (near 1) indicates strong suppressive power. Population suppression with any gene drive strategy requires reasonably high genetic loads. Type and generation rate of resistance alleles, drive conversion efficiency, and fitness costs are the factors that impact the genetic load of a gene drive system^17^. Current CRISPR homing suppression drive systems commonly use haplosufficient female-specific lethal/sterile genes as the target because these will allow for higher genetic load than genes affecting both sexes. The removal of nonfunctional resistance alleles does not readily occur since they can be protected when combined with a wild-type allele^12,18–20^. Instead, they can persist, removing drive alleles in females and preventing drive conversion in males. Consequently, the genetic load is reduced, especially if the drive also has fitness costs. This is a common issue for current homing suppression drives targeting haplosufficient sites, and it diminishes their suppression power.

*doublesex* (*dsx*) has emerged as a good target for homing suppression drive because it is a highly conserved gene in the sex determination pathway of dipterans^21,22^. The regulation of *dsx* relies on sex-specific alternative splicing^23–25^. Mutations in the female-specific exon can lead to dominant female sterility^26^. The female exon of *dsx* can be an ideal target to minimize the formation of functional resistance alleles^19,27–30^. *dsx*-targeting homing suppression drive systems have been designed and tested in *Anopheles gambiae*^19^, *Anopheles stephensi*^29^, *Drosophila melanogaster*^30^, and *Drosophila suzukii*^28^. Mutations at the target site generated nonfunctional resistance alleles in these drives.

However, the discovery of dominant female-sterile resistance in *dsx* from previous studies inspires a promising approach to develop an enhanced homing suppression drive^30,31^. This drive can theoretically exhibit robust suppressive power, enabling the rapid elimination of target populations. Dominant female-sterile mutations were consistently created by three gRNAs targeting the female-specific splicing site of *dsx*, which disrupts original splicing mechanism, leading to the production of male *dsx* transcripts in females and ultimately causing sterility^30^. Once the mutation was generated and transmitted to females, it was be quickly removed through dominant sterility. This resulted in a highly effective self-limiting gene drive, which required far fewer releases of transgenic males compared to traditional pest control strategies such as sterile insect technique. Its advantages can be attributed to its effect on rapidly sterilizing females with resistance alleles. Some recent studies also noticed that dominant mutations can induce a large genetic load on the population, boosting self-limiting suppression, though the pest control system they proposed were not based on CRISPR homing^31,32^.

Here, we propose an improved self-sustaining homing suppression drive by generating dominant female-sterile resistance (named as HSD-Dominant resistance system). We utilized the same gRNA target sites as our previous study^30^. To achieve high drive transmission efficiency as a self-sustaining drive, it is necessary for drive heterozygous females to remain fertile. We used two strategies to avoid expression of male transcripts in females. One was to introduce the strong splicing acceptor site to force drive transcripts to splice to exon 4^33,34^. The other strategy was to restore the original female splicing by using a *dsx* 3′ UTR derived from other *Drosophila* species. However, drive homozygous males were sterile in both these strategies. Thus, several designs were employed to rescue the male transcripts of *dsx* in males by introducing additional sex-specific regulatory elements. Unfortunately, none of those designs rescued fertility of homozygous males, which somewhat reduces overall system effectiveness. However, modeling confirms that both the ideal HSD-Dominant resistance system and the homozygous male-sterile HSD-Dominant resistance system possess substantially greater suppressive power than current “standard” female sterile homing suppression drives that produce only recessive sterile resistance alleles.

## Methods

### Plasmid construction

Our plasmid design was based on TTTgRNAtRNAi, TTTgRNAt, and HSDRed3g, which were constructed previously^12,30^. Original *dsx* fragments were amplified from the genome of *w^11^*^18^ flies. Recoded *dsx* fragments were synthesized by BGI Company. The gRNA target sites were the same as our previous study^30^. Reagents for restriction digest, PCR, Gibson assembly, and plasmid miniprep were obtained from New England Biolabs and Vazyme. PCR primers were from Integrated DNA

Technologies Company and BGI. 5-α competent *Escherichia coli* were from Vazyme. The ZymoPure Midiprep kit from Zymo Research was used to generate materials for injection mixes. Plasmid construction was confirmed by Sanger sequencing. Detailed information about DNA fragments, plasmids, primers, and restriction enzymes used for cloning of each construct are listed in the Supplementary Material, and annotated sequences are available on GitHub (https://github.com/chenwz22/dsxdrive_supplement/).

### Generation of transgenic lines

Embryo injections were conducted by UniHuaii Transgenic Flies Company. The donor/gRNA plasmid (300 ng/ul) was injected into *w^11^*^18^ flies together with TTChsp70c9^35^ (300 ng/ul) to provide Cas9 for transformation. Flies were housed in modified Cornell standard cornmeal medium (using 10 g agar instead of 8 g per liter, addition of 5 g soy flour, and without the phosphoric acid) in a 25 ℃ incubator on a 14/10-h day/night cycle at 60% humidity. The following transgenic lines were generated, each containing a split drive targeting *dsx*:

*s*: Mhc splicing site, stop codon, p10 3′ UTR-tdTomato-3xp3 promotor (reverse), U6:3 promotor- gRNAs.

*sd*: Mhc splicing site, degron degradation tag, stop codon, p10 3′ UTR-tdTomato-3xp3 promotor (reverse), U6:3 promotor-gRNAs.

*sp*: Mhc splicing site, PEST degradation tag, stop codon, p10 3′ UTR-tdTomato-3xp3 promotor (reverse), U6:3 promotor-gRNAs. Lines *sp1* and *sp2* are sublines.

*mA*: *dsx* original splicing site, PEST degradation tag, stop codon, A 3′ UTR (*dsx* female 3′ UTR sequence from *Drosophila innubila* and *melanogaster*), shortened *dsx* intron 4, rescued *dsx* male exon5 and 6, p10 3′ UTR-tdTomato-3xp3 promotor (reverse), U6:3 promotor-gRNAs.

*mB*: *dsx* original splicing site, PEST degradation tag, stop codon, B 3′ UTR (*dsx* female 3′ UTR sequence from *Drosophila melanogaster, suzukii*, and *virilis*), shortened *dsx* intron 4, rescued *dsx* male exon 5 and 6, p10 3′ UTR-tdTomato-3xp3 promotor (reverse), U6:3 promotor-gRNAs.

*C*: *dsx* original splicing site, PEST degradation tag, stop codon, C 3′ UTR (*dsx* female 3′ UTR from other *Drosophila* species), U6:3 promotor-gRNAs, 3xp3 promotor -tdTomato.

*mdsx*: Mhc splicing site, PEST degradation tag, stop codon, SV40-tdTomato-3xp3 promotor (reverse), U6:3 promotor-gRNAs, *dsx* promotor, recoded *dsx* exon 2&3-dsx intron 3, PEST sequence.

*mmsl2*: Mhc splicing site, PEST degradation tag, stop codon, SV40-tdTomato-3xp3 promotor (reverse), U6:3 promotor-gRNAs, *msl2* promotor, recoded *dsx* male CDS, msl2 3′ UTR.

*cctra*: *cctra* intron1 (from *transformer* gene of *Ceratitis capitata*), recoded male exon 5, *cctra* intron 2, PEST sequence, SV40-tdTomato-3xp3 promotor (reverse), U6:3 promotor-gRNAs.

### Phenotypes and morphological analysis

Flies were anesthetized with CO2 and screened for fluorescence using the NIGHTSEA adapter SFA-GR for tdTomato and SFA-RB-GO for EGFP. Fluorescent proteins were driven by the 3xP3 promoter for expression and easy visualization in the white eyes of *w^1118^* flies. tdTomato was used as a marker to indicate the presence of the split drive allele, and EGFP was used to indicate the presence of the supporting Cas9 allele. Morphological photos were taken using a stereo microscope with 10x/22 magnification. To test drive conversion and somatic expression, our drive line was combined with split Cas9 lines (SNc9NG: Cas9 with *nanos* promoter, 5′ UTR, 3′ UTR, and some DNA downstream of the 3′ UTR).

### Fertility and viability assay

Four different cross schemes were conducted to investigate the fertility of male and female drive carriers, as well as the impact of the Cas9 source on fertility (Fig. S1). We first tested the fitness of drive individuals without Cas9 by crossing *w^1118^* males with females from the drive lines *s/sd/sp/Δsp*, which were the parents of the tested and control flies. After 14 days, we collected non-drive flies (without fluorescent eyes) as control individuals, and drive flies (with red fluorescent eyes) as test group individuals. After three days of maturation, each fly was placed in one vial to mate with one *w^1118^* fly of the opposite sex. Every 22 hours, the flies were transferred to a new vial, and we counted the number of eggs in the previous vial. We repeated these steps for three or four iterations, retaining all the vials. We compared the egg number of the test and control groups to assess the reproductive capacity of tested females and males. About 14 days later after all viable offspring had eclosed, we performed phenotypic identification for offspring in all the vials. The offspring survival rate can be calculated based on the number of eggs that developed into adults.

To further understand how the Cas9 source affect the fitness of the drive individuals, we conducted fertility assays on individuals with either "maternal Cas9," "paternal Cas9," or "biparental Cas9" (Fig. S1). For maternal Cas9 test, we crossed *nanos*-Cas9 homozygous females with heterozygous drive males from line *sp1* and *sp2*. For paternal Cas9 test, we crossed *nanos*-Cas9 homozygous males with heterozygous drive females from line *sp1* and *sp2*. For the biparental Cas9 test, we crossed males carrying a heterozygous drive (line *sp)* and homozygous *nanos*-Cas9 with females carrying a heterozygous drive. The progeny of these crosses were then used for drive and fertility experiments as described above.

### Phenotype data analysis

To account for the batch effects (each individual cross is considered as a separate batch with different parameters, which could bias rate and error estimates), we analyzed our data as in previous studies^11,16,36^. In brief, fitting a generalized linear mixed-effects model with a binomial distribution (maximum likelihood, Adaptive Gauss-Hermite Quadrature, nAGQ = 25) enables variance between batches, which then results in marginally different parameter estimates but higher standard error estimates. This analysis was performed with R (3.6.1) and supported by packages lme4 (1.1-21) and emmeans (1.4.2). In our study, these rate estimates and errors were close to the pooled analysis, indicating only minor batch effects at most.

### Diagnostic PCR

To assess transcription of *dsx* in drive and resistance allele carriers, flies were frozen and homogenized. RNA was extracted by using an RNeasy Mini Kit, and reverse transcription was used to obtain cDNA with RevertAid First Strand cDNA Synthesis Kit with oligo(dT) primers. This cDNA was the template for PCR using Q5 DNA Polymerase from New England Biolabs with the manufacturer’s protocol.

Primers Exon3_S_F and tdTomato_S_R were designed to specifically amplify the drive female transcript, and Exon3_S_F and Exon5_S_R were designed to amplify the male-specific dsx transcript.

### Population modeling

Stochastic simulations were performed in SLiM (version 4.0)^37^ similarly to previous studies^11,30,38^. Our simulations have a single panmictic population of generic diploids with discrete generations. The population is defined by the numbers of male and female adults of each genotype. In each generation, each adult female randomly selects a mate from the adult male population and then produces the next generation. To account for crowding and competition typical of most insect systems, we introduce a fitness-based formula for female fecundity (indicated as p in the following formulas). The number of offspring is drawn from a binomial distribution ranging from 0 to 50, with an average value determined by the female’s fitness:

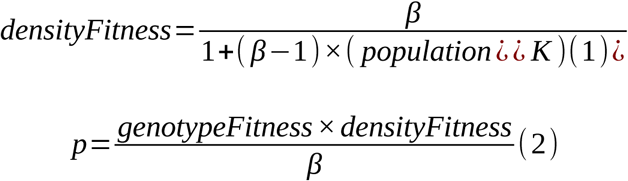

Here, K is the expected carrying capacity and β is the low-density growth rate. Each offspring generated is assigned a random sex, and its genotype is determined by randomly selecting one allele at each genetic locus from each parent, with adjustments for drive activity.

Our model includes wild-type alleles, drive alleles, and (nonfunctional) resistance alleles. We assume that conserved targets sites and multiple gRNAs mitigates functional resistance. For the HSD-recessive resistance system, both the drive allele and resistance allele can induce recessive female sterility. For the HSD-dominant resistance system, the drive allele causes recessive female sterility, whereas the resistance allele leads to dominant female sterility. In one variant of the HSD-dominant resistance system, the drive allele also results in recessive males sterility (resistance alleles do not).

We simulated a complete drive, including both Cas9 and gRNA. Drive/wild-type heterozygous males will convert a fraction of wild-type alleles in their germline into drive alleles at the drive conversion rate. Remaining wild-type alleles are all converted to resistance alleles unless otherwise specified. To account for somatic expression in female drive carriers, a 20% fitness cost was applied to drive heterozygous females unless otherwise specified. Transgenic individuals carry one drive allele and are released after allowing the simulation to equilibrate for ten generations.

Genetic load is defined here as the average net fitness reduction relative to a wild-type population of the same size after the drive reaches an equilibrium. Here, we determine the ratio of actual female population size to the expected female population size without drive to calculate the genetic load. The expected female population size was inferred from the population size of the last generation, based on the low-density growth rate and capacity.

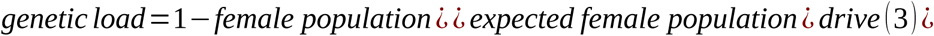

The genetic load for each time step after the drive reaches equilibrium are averaged to find the drive’s genetic load, based on twenty simulation replicates.

## Results

### Dominant resistance improves population suppression

Most current homing suppression drives target haplosufficient female fertility genes, and resistance alleles created by end-joining repair are also recessive female-sterile (we term this system “HSD- Recessive resistance”), assuming that sufficient measures are taken to prevent functional resistance^12,15,36^. However, resistance alleles accumulate in the population, slowing and blocking the spread of drive, thus impeding population elimination. The improved homing drive system we propose here forms dominant female-sterile resistance alleles (“HSD-Dominant resistance”). Once these resistance alleles are transmitted to a female, they will be quickly eliminated by the dominant sterile effect. Thus, the resistance alleles are less able to accumulate in the population compared to recessive sterile alleles, and they contribute directly to female sterilization.

To compare the efficiency of these systems, we simulated them using SLiM individual-based modeling. With a drive that has intermediate performance, our results show that compared to the HSD- Recessive resistance system, the HSD-Dominant resistance system can eliminate the population more efficiently (Fig. 1). The HSD-Recessive resistance system lacks the suppressive power needed for population elimination, so it reaches an equilibrium allele frequency. At this point, the population is reduced but not eliminated.

**Figure 1.**
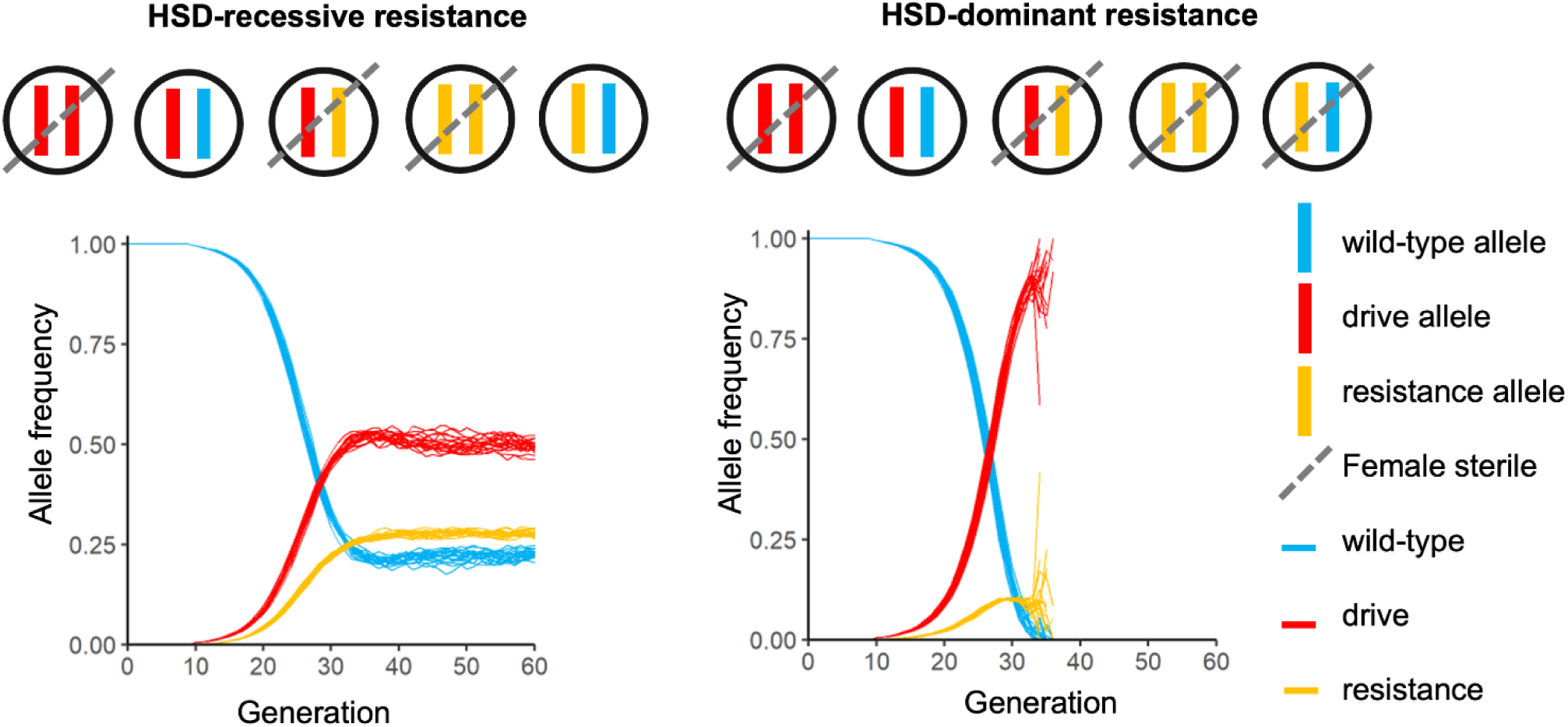
Comparative population dynamics of suppression drives with recessive and dominant resistance. Circles show different genotypes in females, and the dashed line indicates female sterility. The population dynamics simulations are conducted in panmictic populations averaging 100,000 individuals with a low-density growth rate of 6 and a linear density-dependent growth curve. Initial homing suppression gene drive introduction is 0.1% of the total population. The drive conversion rate was set to 80%, the germline resistance formation rate was 15%, embryo resistance formation rate was 20%, and the fitness of heterozygote females was 60% compared to wild-type females. 20 simulations are shown for each drive. All the simulations with dominant-sterile resistance ended in population elimination before generation 40.

### HSD-Dominant resistance system with a Mhc splicing acceptor site

Previously, we constructed a self-limiting suppression drive targeting the female-specific exon of *doublesex* in which the drive was dominant sterile, though it could reliably generate dominant female-sterile resistance (∼95%)^30^. We attempted to represent a self-sustaining (albeit still in split form with Cas9 provided at a distant genomic site) HSD-Dominant resistance system by starting with our earlier construct and rescuing the fertility of drive female heterozygotes. The *dsx* gene is known to usually be a haplosufficient gene, and its dominant sterility arises from expression of the male isoform in females^23,24,26^. We aimed to induce the *dsx*-drive transcript in females to splice to exon 4 even when the drive was present. This was accomplished by introducing a strong splicing acceptor site Mhc, as previously undertaken in *Drosophila suzukii* (though this study did not find dominant resistance)^28^. To ensure that the drive splice site did not lead to a functional product, we also included a stop codon (line *s*), generating a nonfunctional female version of the *dsx* transcript rather than a male-version *dsx* transcript. We expected drive heterozygous females (D/+) to be fertile with the support of a functional wild-type *dsx* transcript. We also included tdTomato driven by the 3xP3 promoter as a fluorescent marker to indicate the presence of a drive allele. To facilitate the *dsx* transcript degradation (thus ensuring that shortened transcript products could not rescue or interfere with *dsx* function), we introduced either a degron (line *sd*) or PEST (line *sp*) protein degradation tag in front of the stop-codon (Fig. 2a). Line *sp1* and *sp2* are sublines of line *sp,* with same construct but different performance (see below). The Cas9 element, required for drive activity, is placed on chromosome 2R and provided through a separate line. It is driven by the germline specific promotor *nanos* and contains EGFP with the 3xP3 promoter^39,40^ (Fig. 2b).

**Figure 2.**
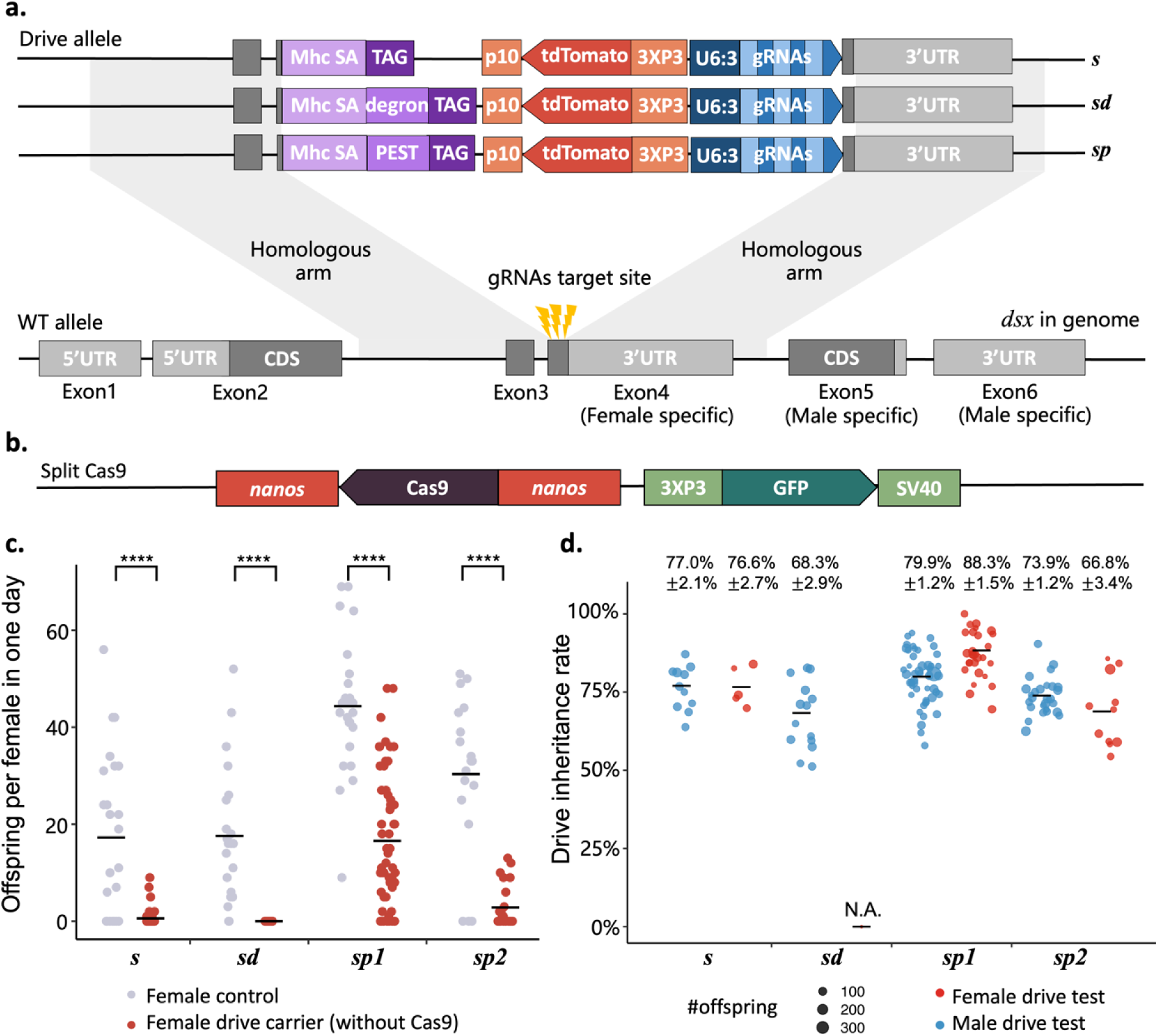
Design and performance of the HSD-Dominant resistance system. **(a)** An Mhc splice acceptor serves as a strong splicing acceptor site, with two additional nucleotides added after the splice site to ensure that the stop codon is in-frame with the rest of the *dsx* gene, thus creating a null copy of the gene. To ensure rapid degradation of the partial *dsx* transcript, one of the constructs has a degron, and one has a PEST sequence. **(b)** The Cas9 element is separately placed on chromosome 2R, regulated by *nanos* promoter/5′ UTR and 3′ UTR. **(c)** Fertility of five drive lines (without Cas9). Each point indicates the number offspring produced by one female during one day, which represents the effects of fecundity and egg viability. Control individuals and tested drive carrier individuals are siblings from the same parent. **** indicates *p*-value <0.0001, t test. Lines *sp1* and *sp2* are sublines of *sp*. **(d)** Inheritance rate of the drive (gRNA) allele. Each point represents the offspring of a single pair of parents. N.A. indicates no offspring for all 16 pairs of parents.

We successfully rescued the fertility of drive heterozygous female in lines *sp1*, *sp2*, and *s* (Fig. 2c). To test drive performance, we first crossed *dsx*-drive heterozygotes with *nanos*-Cas9 homozygotes. The offspring with both green and red fluorescent eyes, indicating that they were heterozygous for the drive allele and Cas9 allele, which were then reciprocally crossed to *w^1118^* of the opposite sex, and their progeny were screened for inheritance of the drive allele (Fig. 2d, Data Set 1 & 2). The drive females of line *sd* were found to be sterile, while drive females of line *sp* were more fertile than those from line *s* (Fig. 2c, Data Set 3). The PEST degradation tag may accelerate degradation of nonfunctional *dsx* transcript, potentially removing an interfering and nonfunctional product and thus accounting for the higher fertility of this line. Line *sp1* was found to cause the highest inheritance bias (80-88%), followed by line *s* (76-78%), line *sp2* (67-74%), and line *sd* (68%) (Fig. 2d). The drive inheritance rates of all tested lines were significantly higher than the 50% Mendelian expectation (*p*<0.0001, z test).

The proportion of intersex individuals among all non-drive female progeny from male drive carriers was moderate, ranging from 10% to 65%, lower than the desired 100% for optimal drive efficiency. This was likely due to a low cut rate (Fig. S2). We performed genotyping on non-drive progeny from male drive carriers of line *s* (Table S1). Sequencing of two intersex females revealed deletions at the splice acceptor site (gRNA target site 1), demonstrating that disruption of this site results in dominant female sterility. Among eight males, three were fully wild-type, one carried a mutation at gRNA target site 3, and four had deletions at the splice acceptor site, which could potentially represent dominant resistance alleles. Among seven phenotypically normal females, two were fully wild-type, three had a deletion at gRNA target site 3, and two carried deletions at both gRNA target sites 1 and 3, but retained an intact splicing acceptor site, representing a recessive resistance allele. Compared to our previous study targeting *dsx*, the dominant resistance formation rate declined, probably because the reduced fraction of simultaneous gRNA cuts and large deletions^30^. It is possible that changing the fluorescent protein (tdTomato is a dimer) or other new drive elements reduced gRNA expression. In contrast, the intersex ratios among the progeny of female drive carriers were significantly higher (ranging from 86% to 100%), particularly when individuals carried the drive allele, suggesting high embryo cutting by maternally deposited Cas9 (Fig. S2), though moderate levels of mosaic cleavage could also potentially have led to this result.

### Fertility of HSD-Dominant resistance system

To further assess fitness costs in drive individuals, we conducted single-pair crosses to evaluate fecundity and fertility. We first examined the impact of drive allele itself. We crossed drive heterozygous males and *w^1118^* females (Fig. S1, Cross Scheme 1). Some progeny from this cross carried one copy of the drive allele, while control individuals were sibling non-drive flies. The number of offspring per day per female represents fertility in our crosses (Fig. 2c). The fertility of line *s* (0.58±0.26, n=48), line *sd* (0±0, n=48), and line *sp2* (2.8±0.9, n=24) was extremely low. Line *sp1* (16.6±1.8, n=57) had a relatively high fertility, but still significantly lower than the control group (*p*<0.0001, t-test). To understand why female fertility may be reduced, we conducted PCR for male-specific *dsx* transcripts, female-specific *dsx* transcripts, and drive transcripts (Fig. S3a). Male transcripts were expressed in the drive females of line *sd*, accounting for its dominant female sterility. Interestingly, the detected male transcript actually included 15-base pairs from the degron of the drive construct, indicating that a splicing donor site may be found in the degron tag (Fig. S3b). Though the expected splicing was still detected in drive individuals (exon 3 followed by exon 4), the unusual male transcript generated by alternative splicing may have been enough to sterilize the female. Note that this unusual male transcript was expressed in both drive males and females of line *sd*. No male transcript was detected in drive females of lines *s*, *sp1* and *sp*2 (Fig. S3a, Fig. 2d). However, drive females with the PEST tag (line *sp1*) nonetheless had higher fertility compared to the drive females without any degradation tag (line *s*). We infer that the PEST protein degradation tag was potentially effective at inducing rapid degradation of the truncated *dsx* protein, thereby reducing its potential impact on the sex development pathway. However, the fertility of drive females was still negatively affected by the drive allele and was significantly lower than the control group (Fig. 2d).

As a split drive, the drive will only be active when combined with Cas9. The source of Cas9 for drive can be from the mother or father (Fig. S1, Cross Scheme 2 & 3). To figure out how the Cas9 source affects the fitness of drive individuals, we conducted fertility assays on individuals that received their Cas9 allele from either a maternal or paternal source for line *sp1* (Fig. 3 and Fig. S4, S5, Data Set 4). Interestingly, the source of Cas9 influenced the fertility and fecundity of the drive females but not males. The drive females with maternal Cas9 had lower fertility than drive females that received their Cas9 allele from their father (Fig. 3). It can be inferred that maternally deposited Cas9 in the embryo, coupled with newly expressed gRNA (which could not have been maternally deposited), may have led to mosaic Cas9 cleavage in late embryonic or somatic cells, potentially negatively affecting sex development and resulting in a significant fitness cost. Note that for line *sp2*, the fertility of drive females remained similar regardless of the Cas9 source (Fig. S5), with the drive allele itself sufficient to nearly eliminate fertility. We also tested drive heterozygotes with homozygous Cas9 (Fig. S1, Cross Scheme 4), which had both maternal and paternal Cas9 alleles. The female fertility reduction was similar to the maternal Cas9 group, likely due to the same amount of maternal Cas9 (Fig. 3).

**Figure 3.**
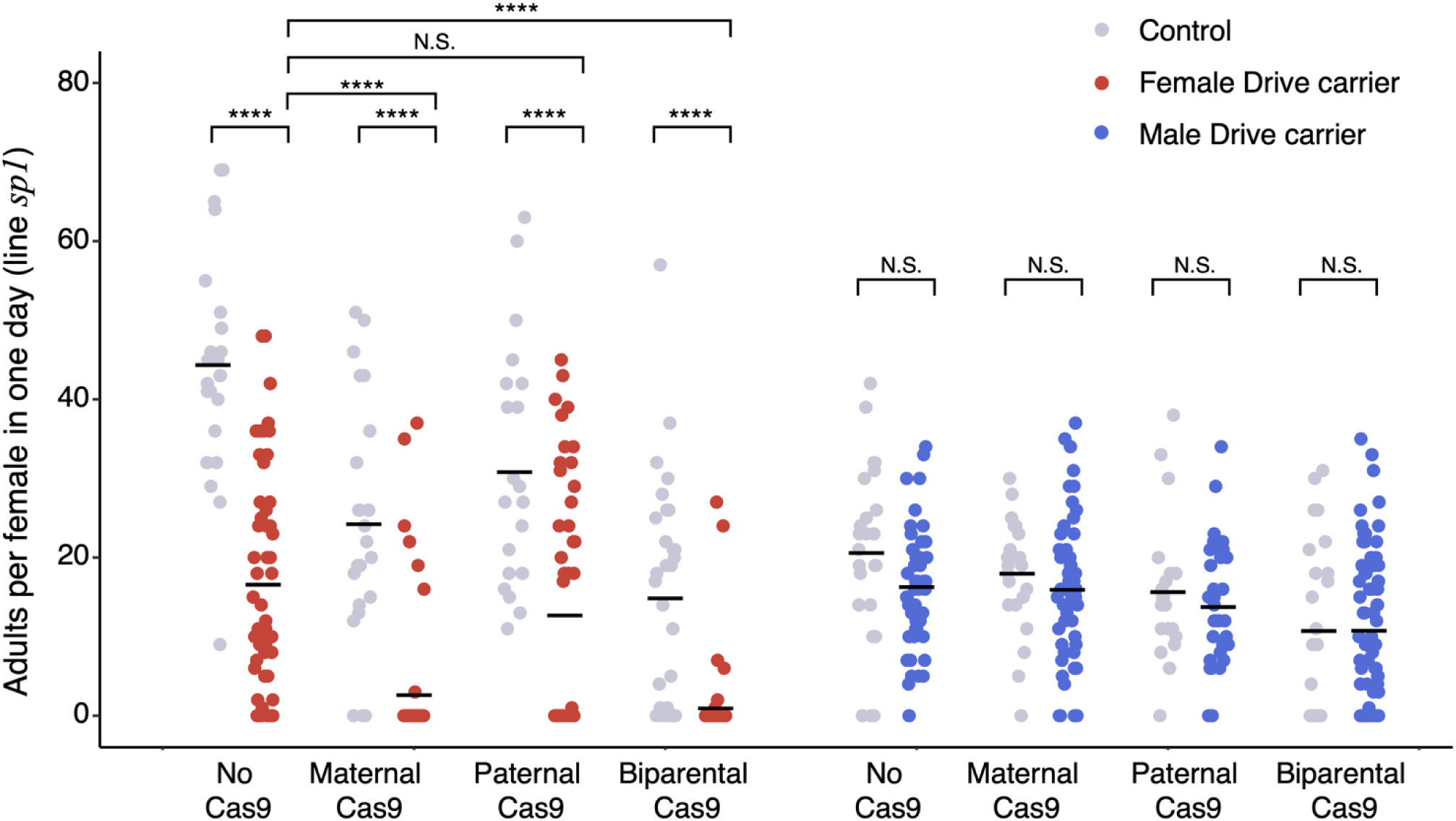
Fertility of the HSD-Dominant resistance system. Four cross schemes were applied to assess the fertility of both males and females from line *sp1*. All the control and tested individuals were siblings with the same number of Cas9 alleles. For the no Cas9 group, parents were drive heterozygous males and *w^1118^* females. For maternal and paternal Cas9 groups (with homozygous Cas9 parents), the other parent was a drive heterozygote. For the biparental Cas9 group, both parents were homozygous for Cas9, and the male was heterozygous for the drive. The t-test was used for statistical comparisons, **** indicates *p* < 0.0001, N.S. indicates no significant difference.

### Sterility of homozygous drive males

Drive homozygotes are expected to exhibit fertility defects in these designs because a strong splice site was used to force splicing to exon 4. This alteration prevents the production of functional *dsx* transcripts, thereby disrupting sexual development in both homozygous drive males and females^28^. To further investigate drive homozygotes, we crossed parents with one copy of both drive ( *line sp*) and Cas9. The abnormal genitalia of intersex individuals cannot be easily classified as male-like or female-like (Fig. 4a). If only drive homozygous females exhibit intersex characteristics, then the ratio of intersex individuals among all drive carrier progeny should be 1/2, assuming relatively high cut rates in the germline and/or embryo, or somewhat less if some wild-type alleles remain. However, a large proportion of drive individuals exhibited intersex phenotype and displayed sterility (125 of 169, *p* < 0.0001 compared to 50% expectation, binomial test). Thus, homozygous males were also likely intersex. To confirm this, we performed Y-chromosome PCR (detecting the Y-linked *Pp1-Y2* gene), which indicated that 3 of 7 intersex flies were genetically male, while the others were genetically female (Fig. 4b).

**Figure 4.**
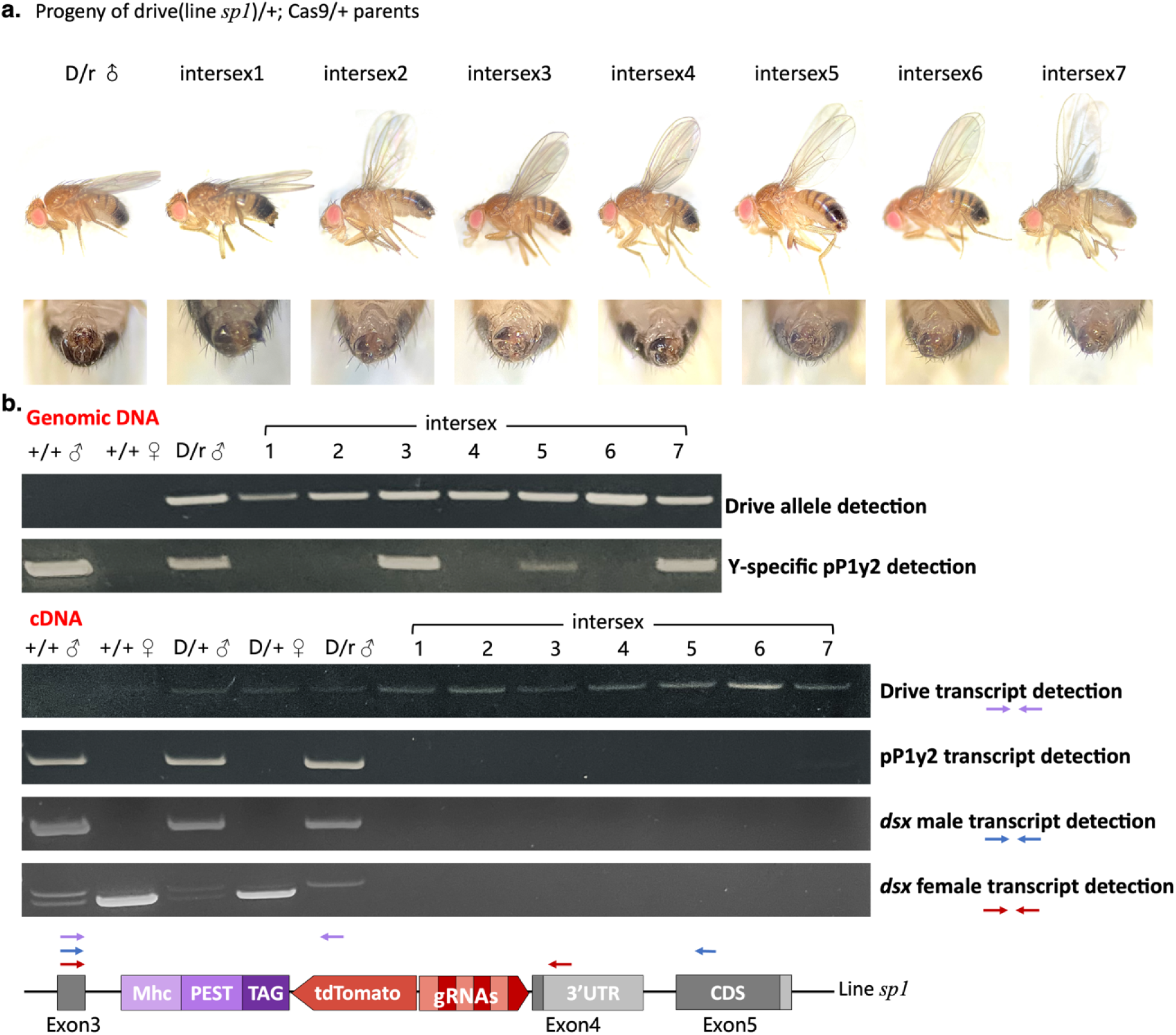
Morphology and *dsx* transcripts in homozygous drive individuals (line *sp1*). **(a)** Morphology of the progeny of parents that were heterozygous for both drive and Cas9. D/r (drive/resistance heterozygotes) males represent progeny exhibiting both male phenotypic characteristics and fertility. All intersex individuals were confirmed to be sterile and were likely mostly drive homozygotes. **(b)** Diagnostic PCR analysis was conducted on both genomic DNA and cDNA of intersex individuals, corresponding to the individuals displayed in (a). +/+ is wild-type, and D/+ is drive heterozygotes. The primer pair exon3_S_F and p10_S_F was used to detect the drive allele (624 bp) and drive transcript (399 bp if intron 3 was spliced). The primer pair pP1y2_F and pP1y2_R was used to detect Y-linked gene pP1Y2, yielding a 591 bp product. The female-specific transcript was detected using primers Exon3_S_F and exon4_S_R, generating a 713 bp amplicon. Similarly, the male-specific transcript was identified with primers Exon3_S_F and exon5_S_R1, producing a 382 bp fragment.

To further understand the splicing mechanism behind the intersex phenotype, we conducted diagnostic PCR to detect *dsx* transcripts by sex-specific primers for these 7 intersex flies. In all intersex flies, neither male nor female transcripts were detected (Fig. 4b). This is attributed to the *dsx*-drive transcript utilizing the strong Mhc splicing site in both males and females, thereby disrupting the normal sex-specific splicing patterns. It is noteworthy that two faint bands were observed in both wild-type males and drive heterozygous males. These bands correspond to the female transcript and a transcript that retained intron 3. The transcript exhibiting intron retention has been identified as *dsx^M2^*, functioning in the nervous system to enhance courtship robustness^28,41^. *dsx^M2^* was also detected in Drive/resistance heterozygous males with deletions at all target sites in the resistance allele. Interestingly, we found that *Pp1-Y2* is regulated either directly or indirectly by *dsx*, as no *Pp1-Y2* transcripts were detected in intersex male individuals (intersex 3,5,7), despite the presence of the gene in their genome. Y-linked *Pp1-Y2* is one of the testis-specific phosphatase genes. It is expressed in the pupa and imago developmental stages and in the testis of males^42,43^. It may be in a downstream pathway of *dsx*, participating in sex development, though a previous study reported that Pp1Y2 knockout would not cause male sterility^44^. Overall, for drive homozygous males, there was no male version of the *dsx* transcript to support correct sex development.

### Alternate HSD-Dominant resistance systems with different *dsx* splicing elements

Because of the Mhc splicing site functions in both sexes, the *dsx* transcript cannot be properly spliced in males. We adopted two general strategies to attempt to recover male splicing, each with three constructs. The first strategy was to preserve the original female splicing acceptor site, followed by several base pairs of CDS region of exon 4, a PEST degradation tag, and an in-frame stop codon (Fig. 5a). Three different *dsx* 3′ UTRs were designed to preserve the highly conserved TRA binding region and termination region^45,46^, but replacing the rest of the 3′ UTR with that from other *Drosophila* species to avoid the risk of partial HDR or reduction in drive efficiency^47^ (Fig. 5a, lines *mA, mB, C*). Considering that the large drive element insertion might change the splicing pattern of *dsx*, a shortened intron 4, male exon 5, and exon 6 of *dsx* were provided in two of these constructs next to the female specific 3′ UTR, aiming to facilitate production of the correct *dsx* male transcript (Fig. 5a, lines *mA, mB*).

**Figure 5.**
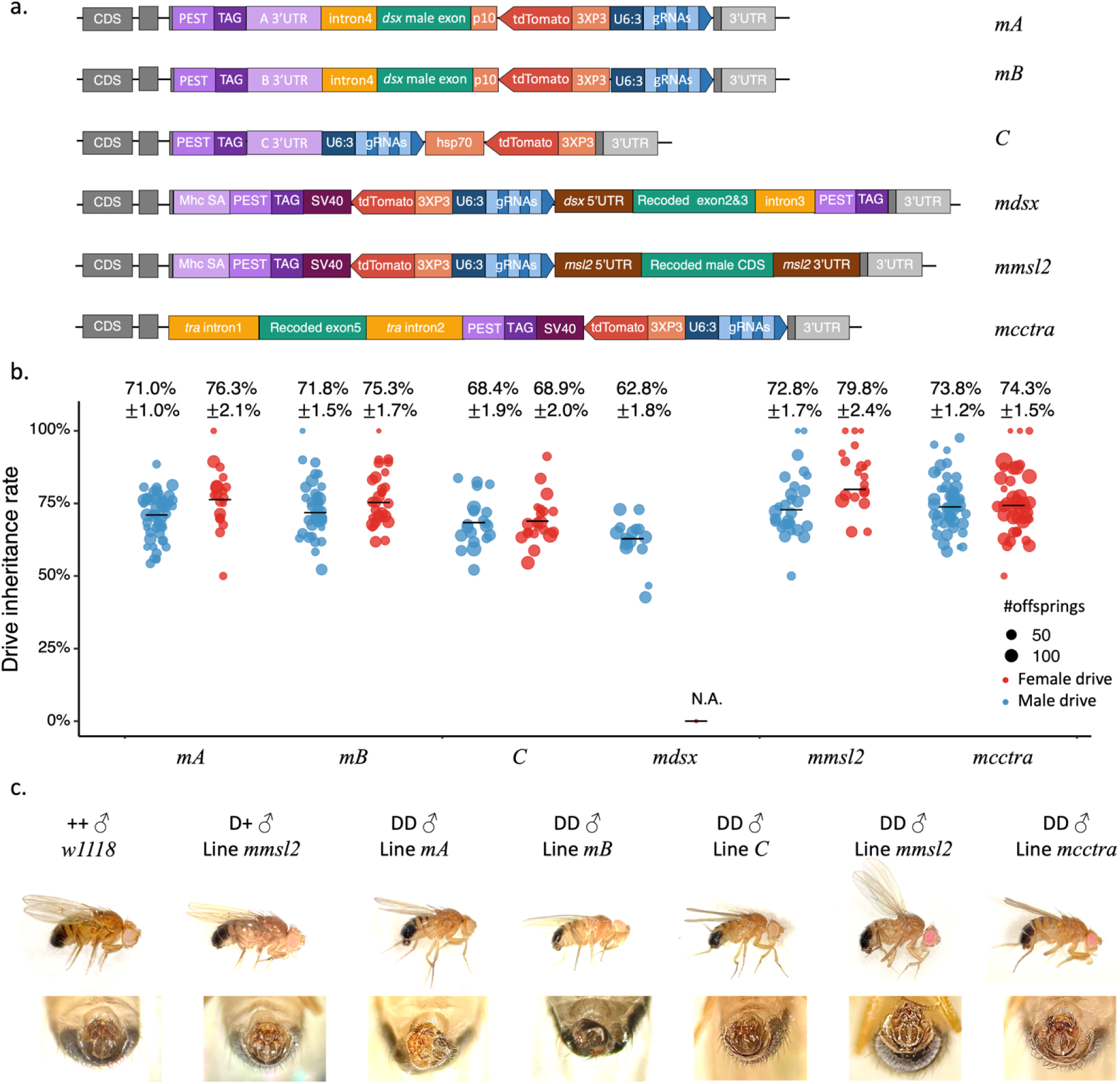
Design and performance of the HSD-Dominant resistance constructs with male rescue. **(a)** The drives of lines *mA, mB*, *and C* preserve the original *dsx* female splice site. The 3′ UTRs of the female exon are derived from other *Drosophila* species. Shortened intron 4 contains 200 bp downstream of *dsx* exon 4 (the female exon) and 200 bp upstream of exon 5 (a male-specific exon). Male-specific exons have the same sequence as the original *dsx* male exons. The left part of the drive construct for lines *mdsx* and *mmsl2* is similar to the construct in line *sp*, but with an SV40 instead of a p10 3′ UTR. *dsx* male rescue was added on the right side of these drive constructs. For line *mcctra*, male-specific *tra* intron 1 is joined with *dsx* intron 3, followed by recoded male exon 5. 3′ *tra* intron 2 is a strong splicing acceptor site. **(b)** Inheritance rate of the drive alleles. Each point represents the offspring of a single pair of parents. **(c)** Morphography of drive homozygous males from lines *mA, mB, C, mcctra*, and *mmsl2*. *w^1118^* males and drive heterozygous males from line *mmsl2* are displayed for comparison.

The second strategy was to introduce other sex-specific regulatory elements to correctly express the *dsx* male transcript (or rescued *dsx* male transcript) in drive males (Fig. 5a). Line *mdsx* terminates the truncated *dsx* transcript on the left side and re-initiates the rescue *dsx* transcript on the right side. We intended that the rescue transcript could utilize the female-specific intron and exon4 in females, while skipping exon 4 and splicing to the downstream exon 5 and exon 6 in males. Line *mmsl2* is designed to terminate the truncated *dsx* transcript and subsequently utilize the male-specific promoter *msl-2* to express a complete *dsx* male transcript in males^48–50^. For line *mcctra,* we joined the *cctra* intron 1 next to *dsx* female-specific exon 3, followed by a recoded *dsx* male exon 5 and exon 6, as well as the *cctra* intron 2. *cctra* intron 1 is male specific, while in females, the transcript will directly splice to the exon after intron 2^51–53^.

Both drive heterozygous males and females were fertile except line *mdsx*, which was dominant female-sterile. The dive inheritance rates of these lines ranged from 68-80%, significantly higher than the Mendelian rate of 50% (*P*<0.0001, z test), but somewhat lower than that of line *sp* (Fig. 5b). We then tested the fertility of homozygous females. For line *mA* and *mB*, we phenotyped the offspring from parents with one copy of both drive and Cas9. We still observed that a large proportion of drive individuals exhibited intersex and displayed sterility (592 of 899 for line *mA*; 732 of 1046 for line *mB*; Data Set 5), which indicates that the homozygous males were also intersex. Y chromosome detection primer pairs were used to confirm the genotype and sex of the intersex flies. 5 of 26 randomly chosen intersex flies among line mA and mB were XY, indicating that intersex flies can be homozygous drive males. For the lines using the second strategy, we phenotyped the progeny from the cross of drive heterozygotes (without Cas9). The fraction of intersex individuals should be 1/6 of all drive carriers in theory if we expect none of the drive homozygous males to be intersex. Interestingly, the observed intersex fractions were 0.164 (n=915), 0.230 (n=437), 0.224 (n=1455) respectively for line *mmsl2*, *mcctra*, and *C* (no statistical significance detected between the observed data and 1/6, binomial test). The morphology of drive homozygous males from these lines closely resembled that of normal males (Fig. 5c). However, follow-up crosses indicate that none of those males were fertile (27 drive homozygous males from line *mmsl2*, 22 males from line *mcctra* and 32 D/D males from line *C* were tested). This indicates that while rescue of fertility was unsuccessful, at least some aspects of the male phenotype were successfully rescued.

### Modeling shows high suppressive power for drives with dominant female-sterile resistance

Fertility of drive homozygous males is important for maintaining a high drive allele frequency in the population. In order to understand how sterile drive homozygous males affect the spread of drive, we first simulated the HSD-Dominant resistance system with sterile homozygous drive males in SLiM (parameters were otherwise identical to Fig. 1). For all simulations, the HSD-Dominant resistance system could still successfully eliminate the population (Fig. 6a). We then compared the population dynamics of HSD-Dominant resistance and HSD-Recessive resistance systems with or without fertile drive homozygous males (Fig. 6b). When the drive reaches equilibrium, the population size in the HSD-Recessive resistance test is suppressed but not eliminated. The system performs substantially worse with sterile homozygous drive males. However, the HSD-Dominant resistance system can fully eliminate the population even with sterile homozygous drive males, albeit after a longer period of time.

**Figure 6.**
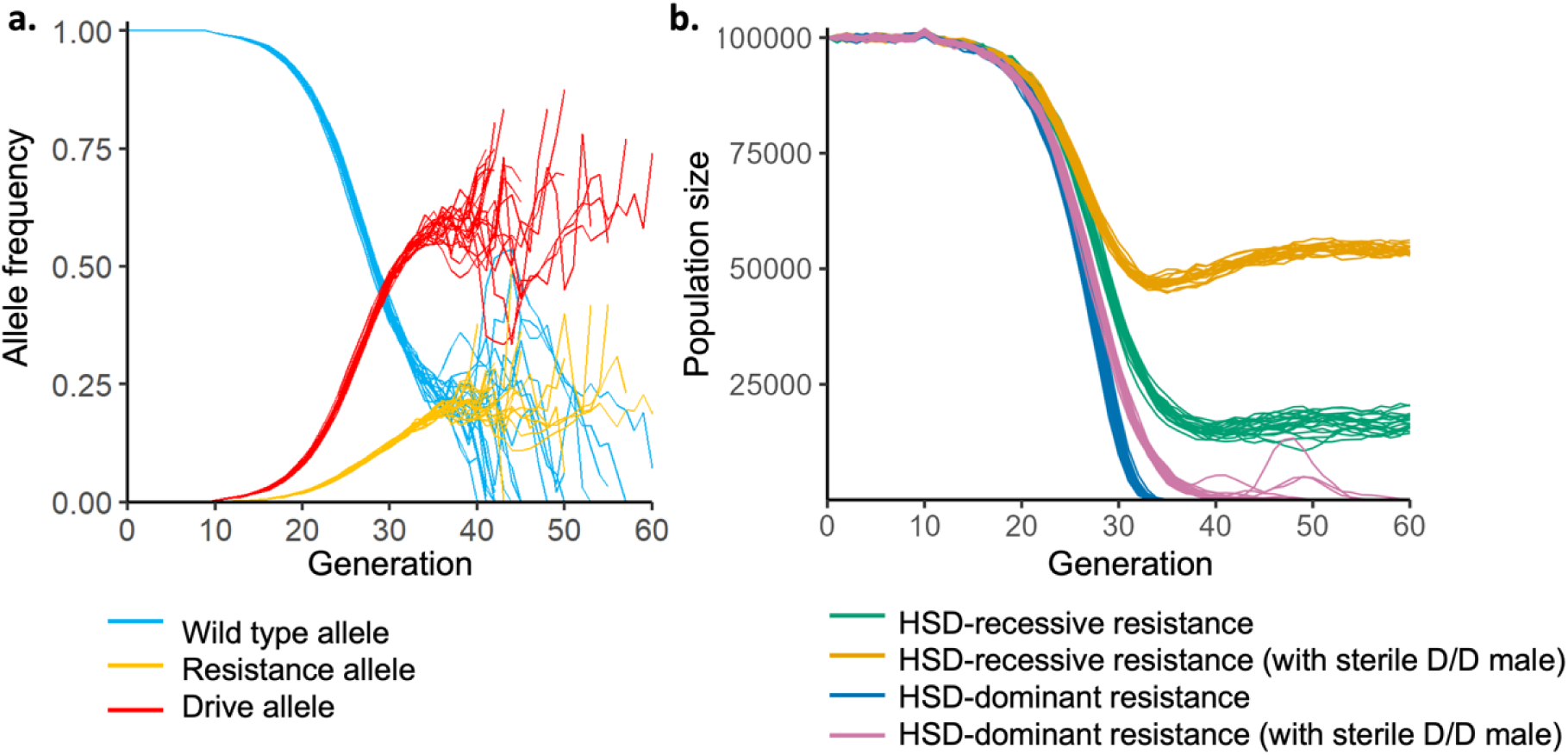
Population dynamics of suppression drives with sterile drive homozygous males. Panmictic population dynamics are simulated with a carrying capacity of 100,000 individuals and a low-density growth rate of 6 with a linear density-dependent growth curve. The initial gene drive introduction is 0.1% of the total population. The drive conversion rate was set to 70%, the germline resistance formation rate was 15%, the embryo resistance formation rate was 10%, and the fitness of heterozygote females was 80% that of wild-type individuals. 20 simulations are shown for each drive. **(a)** Allele frequencies of the HSD-Dominant resistance system with sterile homozygous drive males. **(b)** Population size of each system with varying resistance dominance and homozygous male fertility.

In suppression gene drive analysis, the genetic load is defined as the fractional reduction in the population’s reproductive capacity when drive frequency reaches equilibrium compared to a wild-type population of the same size. It represents a quantitative measure of the suppressive power a drive can achieve. We evaluated the effectiveness of the HSD-Recessive resistance system, HSD-Dominant resistance system, and HSD-Dominant resistance system with sterile drive male homozygotes in a SLiM program framework designed to measure genetic load. We first explored the impact of drive conversion efficiency and germline resistance rate on genetic load (Fig. 7a). No embryo resistance or drive female fitness costs were considered. The HSD-Recessive resistance system requires high drive conversion efficiency to achieve high genetic load, with little effect from germline resistance. HSD-Dominant resistance system can tolerate worse drive conversion and still achieve high total genetic load if the total germline cut rate (drive conversion together with germline resistance formation) is high, which is similar to two-target suppression drives^54^. Even when drive homozygous males are sterile, this remains the case, though genetic load declines more rapidly as the total cut rate falls.

**Figure 7.**
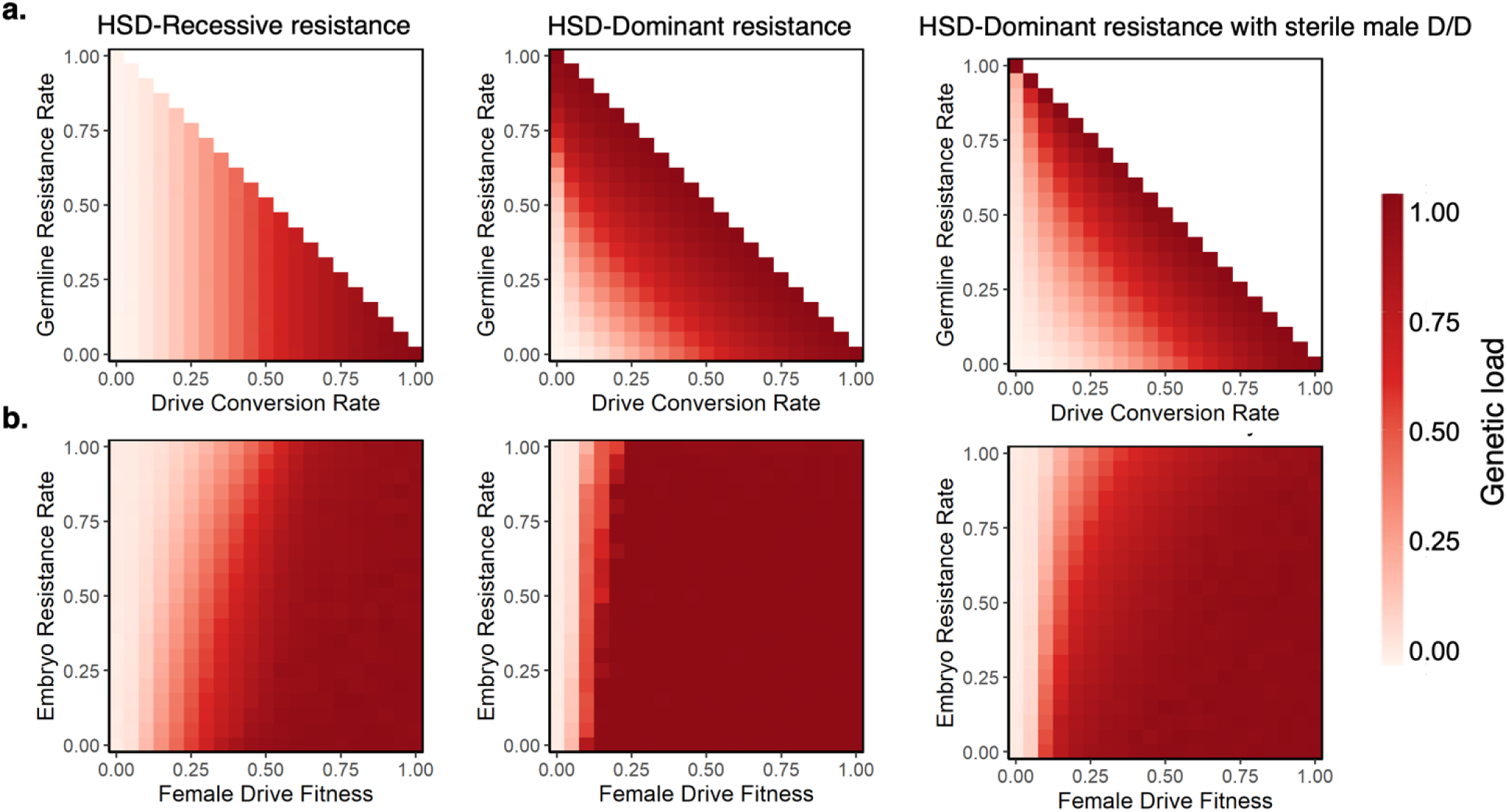
Genetic load of homing suppression drives. Simulations were used to measure genetic load, which represents the suppression power of the drive (reduction in the reproductive output compared to a wild-type population of the same size) when it reaches its equilibrium frequency. **(a)** Genetic load with varying drive conversion and germline resistance, with no fitness costs or embryo resistance. **(b)** Genetic load with varying female drive heterozygote fitness and embryo resistance, with a drive conversion rate of 0.95 and a germline resistance rate of 0.025. Each spot represents the average of 20 simulations.

Embryo resistance and drive female fitness are also key factors influencing the success of a suppression drive. Resistance alleles can be formed in the embryo by maternally deposited Cas9 produced by a drive mother, further hindering drive transmission. High fitness costs in drive females is a common issue in suppression gene drive systems, often caused by leaky expression of Cas9 in somatic cells. If the reproductive ability of drive females is significantly impaired, the efficiency of the drive system will be greatly affected. Thus, we varied embryo resistance rate and drive female fitness with high (Fig. 7B) and intermediate (Fig. S6) drive performance. The HSD-Dominant resistance system can maintain a high genetic load across a wide range of drive female fitness and embryo resistance rate, though it eventually loses its ability to spread. A high performance HSD-Dominant resistance system can even tolerate 100% embryo resistance rate when drive female fitness is as little as 0.25 (Fig. 7B). Sterile homozygous males reduce the genetic load of HSD-Dominant resistance system, but it still is often substantially higher than the HSD-Recessive resistance system.

## Discussion

In this study, we proposed an improved pest control strategy, homing suppression drive generating dominant female-sterile resistance alleles (HSD-Dominant resistance), which we have shown to be feasible and stronger than current homing suppression drive systems that generate recessive sterile resistance alleles (HSD-Recessive resistance). Our demonstration in flies successfully rescued heterozygotes females and generated dominant resistance, but homozygous males were sterile due to disruption of male *dsx* splicing. This reduced suppressive power in models, but it still remained higher than standard suppression drives when total germline cut rate was high.

CRISPR homing drive relies on homology-directed repair. However, end-joining is also a common repair mechanism in species other than *Anopheles* where homing drive has been tested^2,4^. Thus, the emergence of resistance alleles cannot be completely avoided, which may slow the spread of the drive or even lead to its failure. A variety of strategies have been proposed to mitigate the effect of resistance, including using germline-specific Cas9 promotors, using multiple gRNAs, and targeting highly conserved regions of essential genes^11,12,18,19,40,55^. These mostly address functional resistance, and nonfunctional resistance can be perhaps only partially mitigated by improved promoters. Here, inspired by the dominant resistance observed in previous study targeting the female exon of *dsx*, we aimed to construct homing drive system where resistance alleles can be easily eliminated by dominant female sterilty^30^. HSD-Dominant resistance has high genetic load for a large parameter range. The advantages of HSD-Dominant resistance system are stronger when the total germline cut rate is high. Germline resistance contributes to population suppression because resistance alleles in the HSD-Dominant system will quickly be eliminated while reducing reproduction in female carriers. The HSD- Dominant system is also much less sensitive to embryo resistance and female fitness compared to the HSD-Recessive resistance system, though fitness in particular remains important (Fig. 7, S6).

*dsx* is a haplosufficient gene, and its dominant sterility arises from incorrect sex-specific splicing caused by the insertion of the drive element or a mutation at the splicing acceptor site^30^. Our previous homing drive at this site was itself dominant female-sterile, making it a self-limiting system. To design a HSD-Dominant resistance system based on *dsx*, the key was to ensure that drive female heterozygotes were fertile. We tested three strategies to restore the fertility of drive females. The first was to introduce a strong Mhc splicing site, forcing the *dsx*-drive transcript to use the female-specific exon and generating a nonfunctional female-version *dsx* transcript rather than a male-version *dsx* transcript in females. The second was to preserve the original characteristics of *dsx* structure as much as possible. We restored the original *dsx* female specific splicing site and provided a modified *dsx* 3′ UTR derived from other *Drosophila* species. The third was to guide sex-specific splicing by introducing *cctra* introns. Most of them worked and successfully rescued the fertility of drive heterozygous females. A PEST protein degradation tag was useful for accelerating the degradation of nonfunctional *dsx* transcript and appeared to help increase female fertility, though it was not fully restored for unknown reasons. Perhaps *dsx* is partially haploinsufficient, or perhaps other tissue-specific effects were in play. The partial protein from a drive allele may also be harmful, and a PEST sequence may be insufficient to completely stop it from interfering with normal regulation.

Despite our efforts to preserve male fertility, we observed that homozygous drive males were sterile and intersex in all of our constructs where this could be tested. For lines *sp*, *mA*, and *mB*, *dsx*-drive transcripts used the female exon-specific site for splicing in males. Therefore, there were no functional male transcripts to support the sex development of homozygous drive males. For *lines C, mmsl2*, and *mcctra*, homozygous drive males exhibit visual characteristics similar to those of normal males; however, they remain sterile (Fig. 5c). This may be because the expression timing of the rescue male *dsx* is not precisely the same as that of the original *dsx*. There may also be other unknown sequences that play an important role in splicing the native *dsx* gene. This result differs from the *dsx* drive in *Anopheles stephensi* and *gambiae*^19,29^, possibly because the sex development pathway is different. While *dsx* expression in *Drosophila* is regulated by *tra*, this regulatory mechanism appears to be lacking in *Anopheles*^56^. The *A. gambiae* drive did not suffer from male sterility, while the *A. stephensi* drive was able to still express male transcripts despite sterility.

The sterility of drive homozygous males reduces the suppressive power of HSD-Dominant resistance system, especially when embryo resistance is high and female fitness is low. However, even this imperfect HSD-Dominant resistance system still has stronger suppressive power than HSD-Recessive resistance (Fig. 7). In the HSD-Recessive resistance system, sterility of drive homozygous males significantly reduces the suppressive power. Conversely, in the HSD-Dominant resistance system, the sterility of drive homozygous males does not severely affect genetic load if total germline cut rates remain high, albeit a longer time required for population elimination.

Dominant resistance allele formation rate, while not low, nonetheless declined compared to our previous study, likely due to lower germline cut rate^30^. While the reason for this is unclear (none of the changes to the construct would normally be expected to change gRNA expression), this issue can be addressed by optimizing the gRNA cassette. For instance, we could use more efficient tRNAs or ribozymes to express multiple gRNAs, or employ individual promotors for each gRNA^57–59^. Increasing the germline level of Cas9 can also help, as could eliminating identical DNA in the fluorescent marker. Despite the reduced dominant resistance rate, several resistance alleles with remaining wild-type target sites could potentially be cleaved in males when paired with a drive allele in subsequent generations, providing another opportunity for drive conversion or the formation of dominant resistance.

Additionally, we found that the source of Cas9 affects the fitness of drive females (Fig. 3). Previous research reported that the deposition of the Cas9/gRNA complex from the mother induces fitness costs among female progeny^12^, but here, our result shows that deposited Cas9 itself coupling with newly expressed gRNAs can also induce cuts and harm fitness^36^. This indicates that split drive cage experiments may underestimate the performance of complete drive systems if maternal deposition is moderate to high.

Given that the *dsx*-based sex determination pathway is highly conserved among dipterans, including mosquitos, moths, and flies^22,56,60^, our improved homing drive system utilizing *dsx* sex-specific splicing with dominant effects had good prospects for application in the control of a broad range of non-model pest species. Indeed a dominant resistance allele in *dsx* was recently reported in *Anopheles gambiae*^31^, and it may be possible to consistently generate such alleles with a suitable multiplexed gRNA targeting strategy.

Overall, our study demonstrates an improved homing suppression drive system by ensuring that resistance alleles were dominant female-sterile, facilitating their rapid removal and thereby increasing the suppression power of the drive. With 100% germline cutting, genetic load is maximized, even with modest drive conversion efficiency. We showed that this design indeed proved to be stronger than current homing suppression drive designs that target haplosufficient female fertility genes. However, construction of this system could be partially inhibited by sterility of drive homozygous males. Future studies could potentially address this issue, but even with homozygous male sterility, modeling indicates that the system power was still improved. It is thus possible that highly efficient population suppression gene drives can be constructed in a wide variety of non-model species when nonfunctional resistance alleles have a dominant effect.

## Acknowledgements

This study was supported by the Center for Life Sciences and the National Science Foundation of China (grant 32270672). The cluster-based data collection was assisted by High-Performance Computing Platform of the Center for Life Science at Peking University.

## Supplementary Information

**Figure S1.**
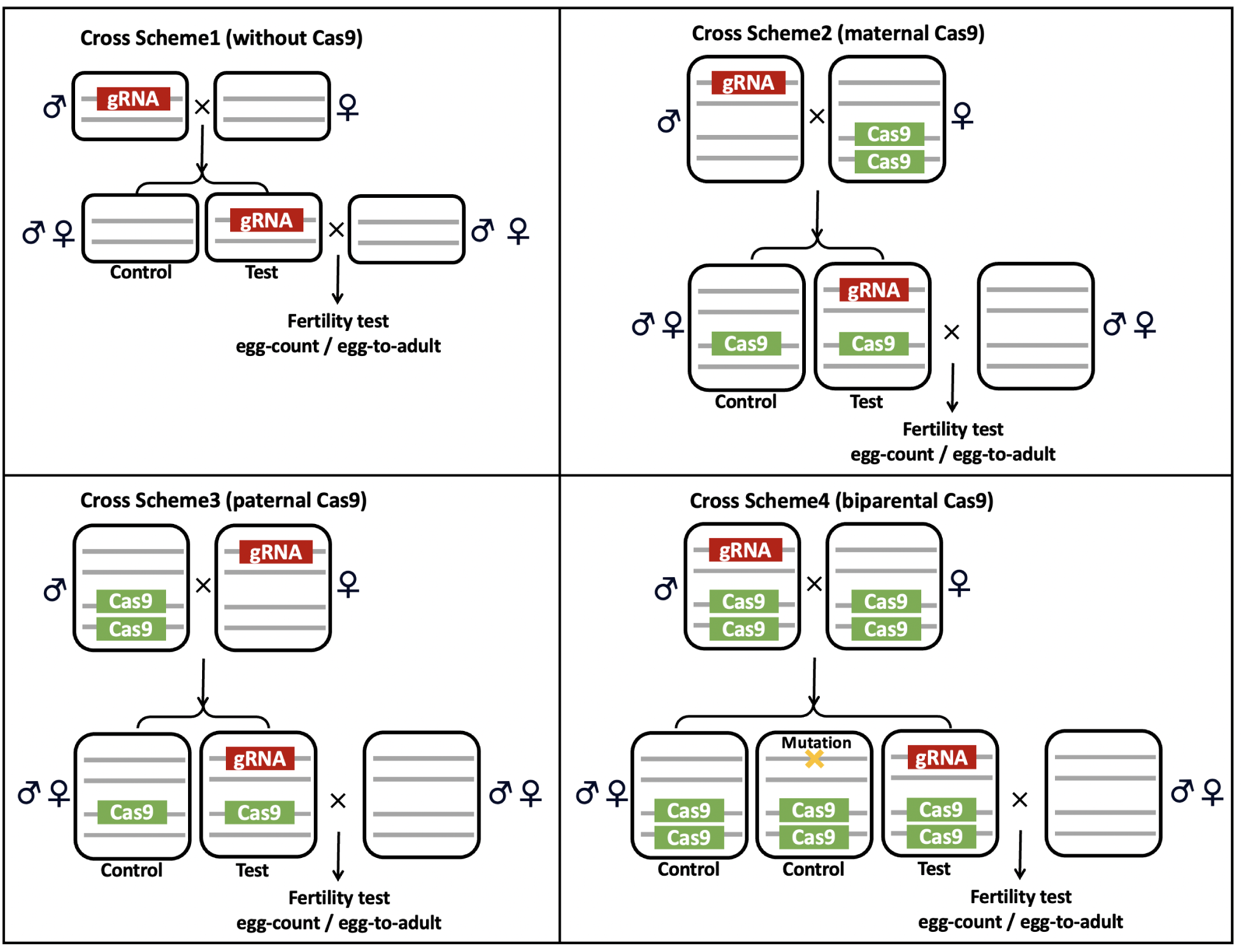
Illustration of cross schemes for fertility tests. Cross scheme 1: no Cas9, which was designed for investigating the impact of drive alleles on fertility. Cross scheme 2 and 3: Cas9 was provided from the father or mother of the tested individuals, in order to assess the effect of maternal or paternal source of Cas9. Cross scheme 4: males heterozygous for the drive and homozygous for Cas9 were crossed to Cas9 homozygous females to generate drive and non-drive flies for fecundity and fertility assessment. The fitness of drive females was likely affected by maternally deposited Cas9 and zygotically expressed gRNA.

**Figure S2.**
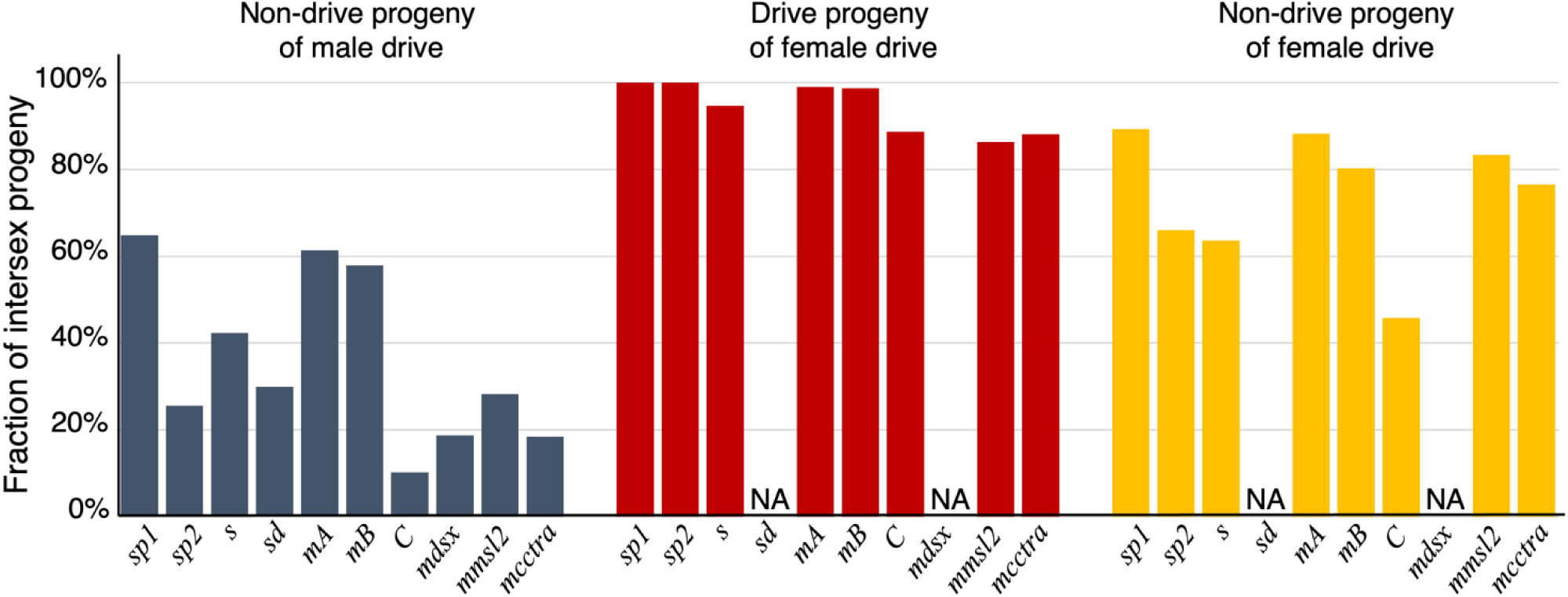
Fraction of intersex females. Drive males or females that were heterozygous for both drive and Cas9 were crossed to *w^1118^* flies, and their offspring were phenotyped. The intersex phenotype was characterized by the presence of a black stripe at the end of the abdomen and in some cases, malformed genitalia.

**Figure S3.**
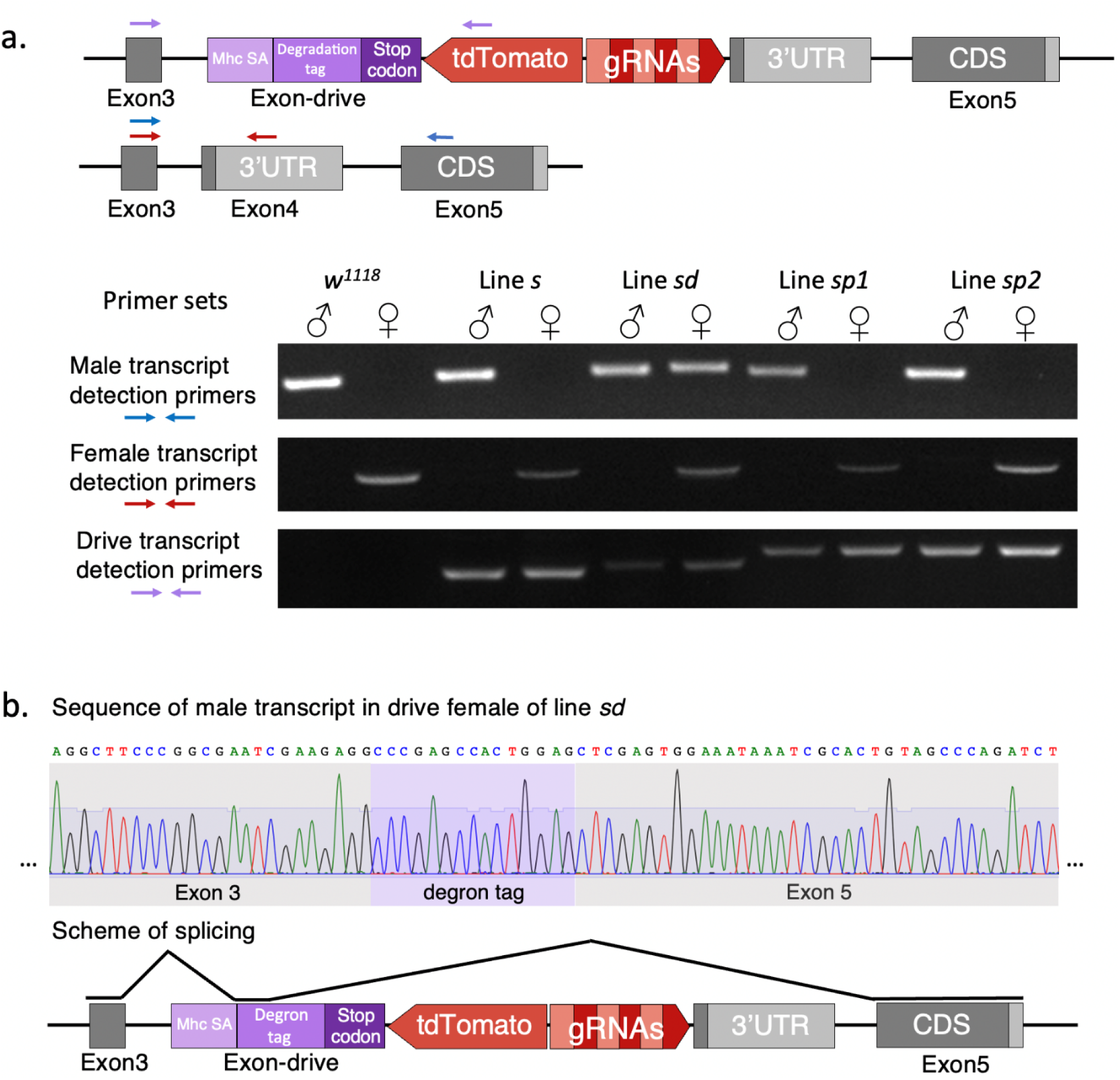
Transcript detection in lines *s, sd, sp1* and *sp2.* **(a)** Three pairs of primers were designed to detect male transcript, female transcript, and drive transcript. The expected amplicon size for the male-specific transcript was 382 base pairs when amplified using primers exon3_S_F and exon5_S_R. The expected amplicon size for the female-specific transcript was 713 base pairs when amplified using primers exon3_S_F and exon4_S_R. The degradation tag is different across different lines, resulting in slight variations in the band length of the drive transcripts. Specifically, the expected amplicons sizes were 341 bp, 350 bp, 401 bp, and 401 bp for lines *s*, *sd*, *sp1*, and *sp2,* respectively. The primers used were exon3_S_F and p10_S_F. **(b)** The sequence of the drive transcript detected in females of line *sd*. The scheme of the splicing mechanism was inferred based on the mRNA sequence.

**Figure S4.**
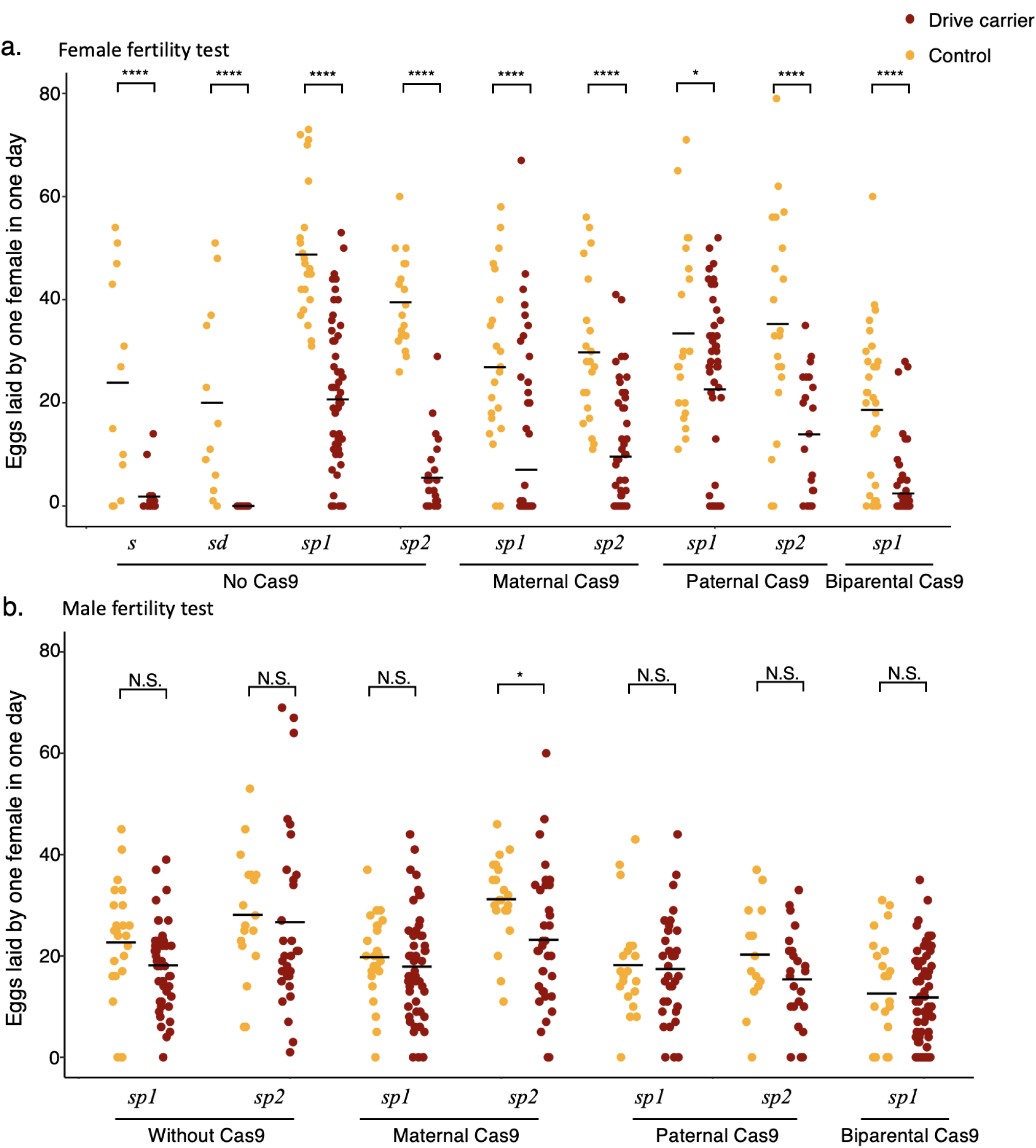
Fecundity of lines *s, sd, sp1,* and *sp2* of the HSD-Dominant resistance system. Number of eggs per day per female was measured to assess fecundity. **(a)** The fecundity of drive females and controls. **(b)** The fecundity of males with their controls. Fecundity assays were conducted as part of fertility assays with varying cross schemes. Control individuals were non-drive siblings of drive carriers. t-tests were conducted for comparisons between drive carriers and controls. N.S. indicates no significant difference. * indicates *p* < 0.05, **** indicates *p* < 0.0001.

**Figure S5.**
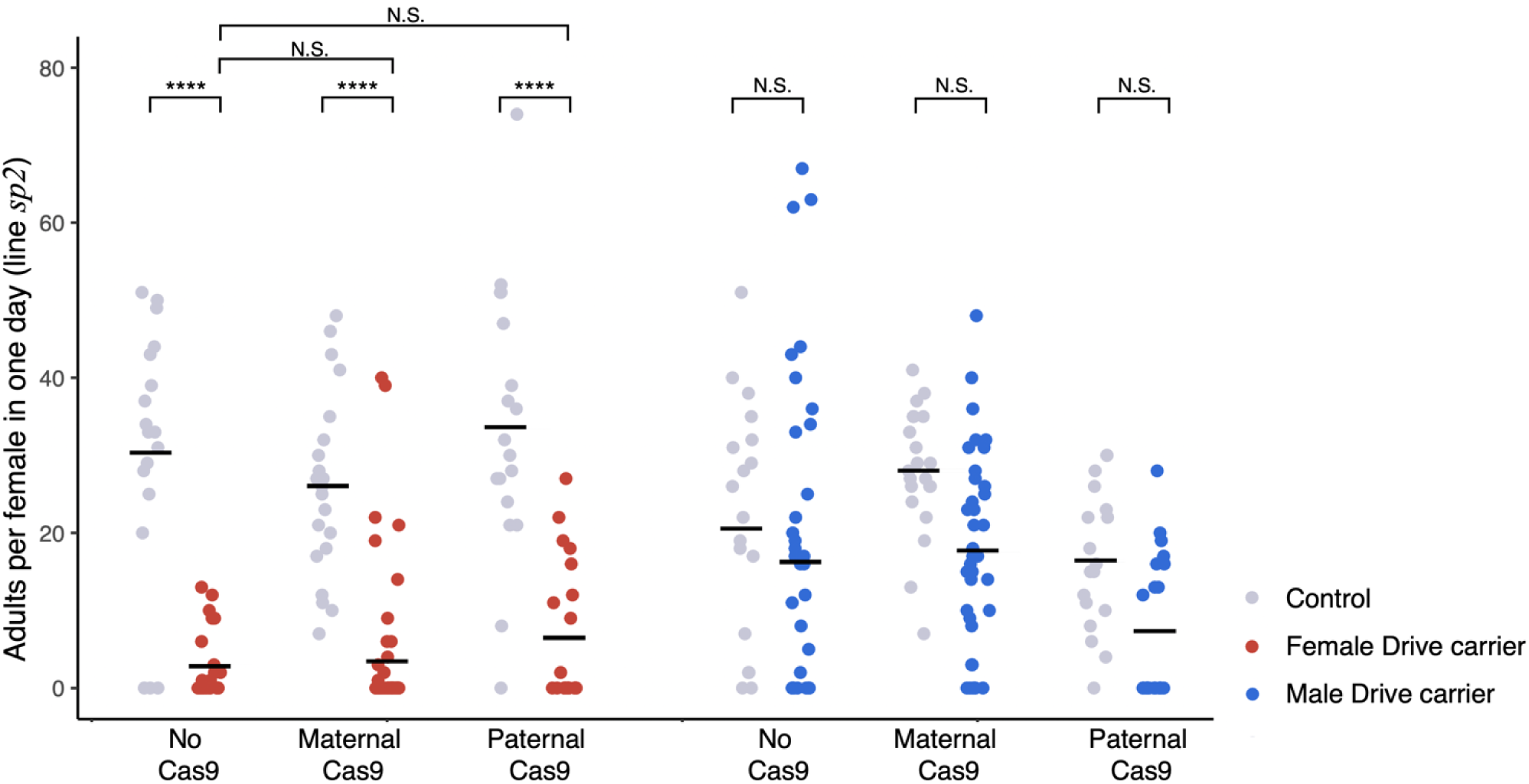
Fertility of the HSD-Dominant resistance system (line *sp2*). Four cross schemes were utilized to assess the fertility of both males and females from line *sp2*. All the control and tested individuals were siblings with the same number of Cas9 alleles. For the no Cas9 group, parents were drive heterozygous males and *w^1118^* females. For maternal and paternal Cas9 groups (with homozygous Cas9 parents), the other parent was a drive heterozygote. The t-test was used for statistical comparisons. **** indicates *p* < 0.0001, N.S. indicates no significant difference.

**Figure S6.**
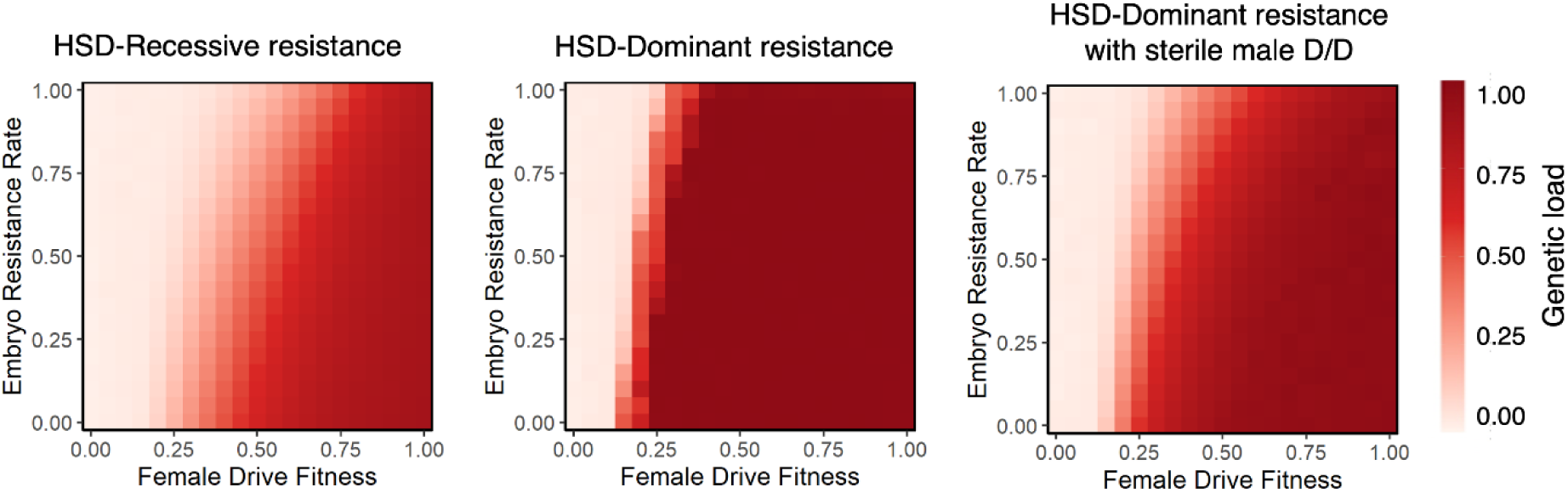
Genetic load comparison with reduced drive performance. Comparison of HSD-Recessive resistance system, HSD-Dominant resistance system, and HSD-Dominant resistance system with sterile homozygous drive males. Genetic load is shown with varying female drive heterozygote fitness and embryo resistance. The drive conversion is 0.8, and the germline resistance rate is 0.15.

**Table S1.** Resistance allele sequences. All sequences are from non-drive progeny of D/+, Cas9/+ fathers and +/+ mothers. WT – wild-type

|  | gRNA target 1 | gRNA target 2 | gRNA target 3 | Splicing acceptor site of exon4 |
| --- | --- | --- | --- | --- |
| Male #1 | cleavage | WT | cleavage | deleted |
| Male #2 | WT | WT | WT | preserved |
| Male #3 | cleavage | cleavage | cleavage | deleted |
| Male #4 | cleavage | WT | cleavage | deleted |
| Male #5 | cleavage | WT | cleavage | deleted |
| Male #6 | WT | WT | cleavage | preserved |
| Male #7 | WT | WT | WT | preserved |
| Male #8 | WT | WT | WT | preserved |
| Female #1 | cleavage | WT | cleavage | preserved |
| Female #2 | WT | WT | WT | preserved |
| Female #3 | WT | WT | cleavage | preserved |
| Female #4 | WT | WT | cleavage | preserved |
| Female #5 | cleavage | WT | cleavage | preserved |
| Female #6 | WT | WT | WT | preserved |
| Female #7 | WT | WT | cleavage | preserved |
| Intersex #1 | cleavage | cleavage | WT | deleted |
| Intersex #2 | cleavage | WT | cleavage | deleted |

